# 3D culture of pancreatic cancer cells *in vitro* recapitulates an aberrant mitochondrial oxidative phosphorylation genotype observed *in vivo*

**DOI:** 10.1101/2024.03.08.583569

**Authors:** Krzysztof Kuś, J. Mark Treherne, David L. Earnshaw, Renata Krzykawska, Beata Biesaga, Małgorzata Statkiewicz, Katarzyna Unrug-Bielawska, Zuzanna Sandowska-Markiewicz, Michal Mikula, Javier Antonio Alfaro, Marcin Przemyslaw Krzykawski

**Affiliations:** Department of Biochemistry, University of Oxford, Oxford, UK; Real Research Sp. z.o.o., 30-348 Kraków, Poland; Talisman Therapeutics Limited, Babraham Research Campus, Cambridge, UK; Department of Genetics, Maria Sklodowska-Curie National Research Institute of Oncology, 02-781 Warsaw, Poland; International Centre for Cancer Vaccine Science, University of Gdańsk, Gdańsk, Poland; Department of Biochemistry and Microbiology, University of Victoria, Victoria, British Columbia, Canada; School of Informatics, University of Edinburgh, Edinburgh, UK

## Abstract

The treatment of many aggressive cancers remains a significant unmet medical need. An aberrant dependence of tumors on mitochondrial oxidative phosphorylation pathways has been well characterized in mediating the progression of pancreatic cancers, as well as some other malignancies. However, the discovery and development of new therapeutic strategies targeting or manipulating this pathway in pancreatic tumors has been relatively slow when compared with other cancers, limiting the current options for treating patients. One key technical challenge in discovering new therapies has been the limited ability of cancer cell lines when grown in 2D conditions to recapitulate the mitochondrial oxidative phosphorylation pathways in vitro that have been observed in vivo. In part, this is because 2D cultures fail to mimic the extracellular 3D tumoral environment. More generally, the translation of data based on 2D pancreatic cancer cell culture systems have also been constraining options for discovering new therapies. Conversely, 3D cultures can provide an improved technology platform in which pathologically relevant pathways in tumors can be recapitulated and analyzed in vitro. In this study, we have demonstrated in 3D cultures the reconstruction of the mitochondrial oxidative phosphorylation genotype in vitro, which more closely resembles that observed in vivo. Our study highlights the value of using transcriptomic readouts as a method to demonstrate the relevance and utility of 3D preclinical models *in vitro*, when grown in a more physiological extracellular environment. This key technological advance can now better enable the discovery and subsequent development of new therapeutic strategies targeting disease-relevant pathways found in pancreatic and other tumors.

## Introduction

Recent decades have seen considerable progress in the diagnosis and more effective treatment of many different cancers but the same cannot be claimed for pancreatic cancers. The most common type of pancreatic cancer is pancreatic ductal adenocarcinoma (PDAC), which is a highly devastating disease with poor prognosis and a rising incidence [1]. Late detection and particularly aggressive biology are the major challenges that underlie the high rates of therapeutic failure. PDAC is currently the fourth most frequent cause of cancer-related death and is projected to become the second most lethal tumor type by the year 2030 [2]. Chemotherapy typically only extends overall survival to about 11 months and very rarely results in long-term and progression-free survival exceeding 5 years [3]. Consequently, innovative approaches are needed to identify novel and more effective therapies for PDAC to overcome resistance to chemotherapy and other more targeted treatments. Currently, some targeted therapies have been approved for specific groups of patients with pancreatic cancer, depending on the presence of specific mutations, copy-number changes or genetic structural variants. Such drugs include erlotinib, olaparib and pembrolizumab [4]. However, such therapies have still not yet made significant breakthroughs to provide real hope for the long-term treatment of PDAC patients. Encouragingly, new research indicates that by targeting the oxidative phosphorylation (OXPHOS) pathway, which has been associated with aberrant metabolism in cancer cells, significant reductions in drug resistance can be observed [5,6]. Current models of pancreatic cancer metabolism acknowledge the coexistence of aerobic glycolysis (or the classical Warburg model) alongside OXPHOS in cancer cells and that metabolic regulation can be flexible, adapting to changing environmental conditions [7]. Evidence also suggests that under conditions of limited access to nutrients, pancreatic cancer cells exhibit a particular dependence on the process of mitochondrial OXPHOS [8]. The therapeutic significance of this process is further highlighted by the ability of cancer cells to withstand low glucose conditions to survive and grow. In cells resistant to low glucose conditions, increased oxygen consumption is often observed compared with more glucose-sensitive cell lines. These observations are consistent with a hypothesis in which cancer cells become well-adapted to challenging metabolic conditions and prioritize the reprogramming of their metabolism to maximize energy efficiency through enhanced mitochondrial respiration. This metabolic shift ensures the production of a sufficient amount of adenosine triphosphate (ATP) to power critical cellular processes [5–9]. This general phenomenon is illustrated in Fig 1.

**Fig 1.**
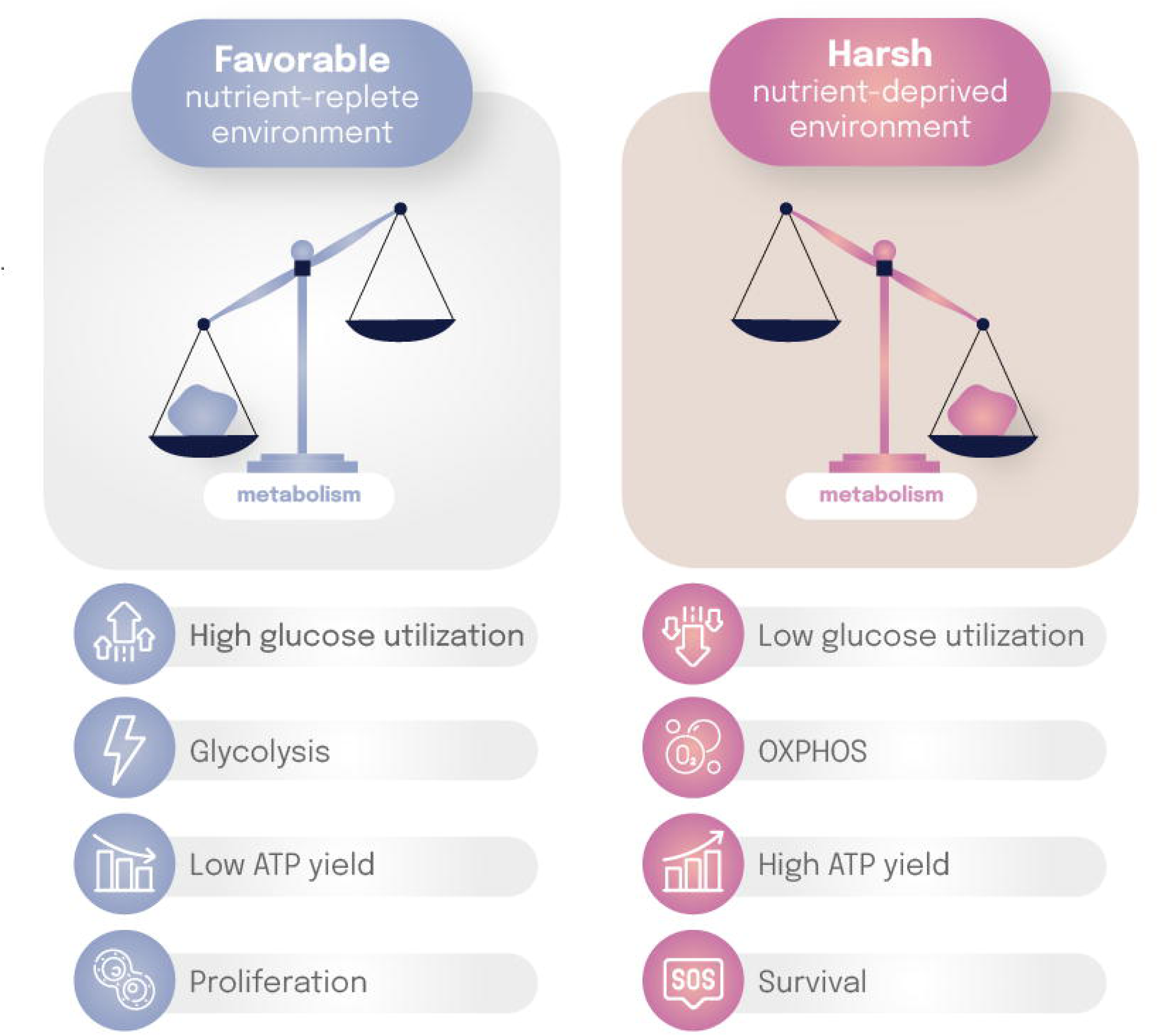
The metabolic dependence of pancreatic cancer cells on the process of mitochondrial oxidative phosphorylation (OXPHOS). PDAC cells have a proliferative phenotype and macromolecular synthesis is prioritized over ATP generation. Under nutrient deprivation conditions, PDAC cells develop a survival phenotype and nutrient conservation to maximize ATP generation is prioritized.

Historically, most preclinical *in vitro* studies used in oncology research have typically relied on the use of conventional ’flat’, 2D or monolayer cultures of cell lines [10]. However, overall success rates in pharmaceutical research have been highly unpredictable, especially for the discovery and development of novel targeted cancer therapeutics [11]. However, more predictive pre-clinical screens are now being more effectively implemented into drug discovery workflows to score therapeutic candidates in a way that better correlates with their potential clinical utility [12,13]. It has been argued that the biological and phenotypic complexity of 3D cultures allows for more predictive assays of future clinical efficacy to be developed, as they are grown in a more physiological and/or pathological 3D environment [14]. In particular, the key role of novel hydrogel scaffolds in mimicking this extracellular environment has been emphasized in transforming the physiological and therapeutic relevance of such 3D cell-based *in vitro* assay systems [15]. The microenvironment for tumor-like growth is of particular importance when testing new drugs, so that the data obtained can be better cross-correlated with *in vivo* studies.

There is emerging evidence that the metabolism of cancer cells cultured in nutrient-rich *in vitro* environments may not fully recapitulate the growth of tumors and 2D culture, which may further obscure metabolic dependencies due to more artificial growth conditions [16,17]. Valle *et al.* developed a novel method of enriching 2D cultures with subpopulations of cancer stem cells [9]. These authors demonstrated the plasticity of PDAC cells maintained in long-term 2D culture conditions with a medium enriched with galactose, which served as an alternative sugar to glucose, whereby the efficiency of ATP synthesis is much lower. Long-term culture with galactose was found to trigger the compensatory activation of OXPHOS pathways. As a result, glycolysis-dependent cells did not survive in the galactose-rich medium, whereas cells with an efficient OXPHOS system survived and became enriched in those cultures. Surviving cells expressed multiple cancer stem cell biomarkers, overexpressed pluripotency-associated genes and related pathways, demonstrated increased promoter activity, enhanced self-renewal capacities and exhibited significantly higher tumorigenicity when transplanted *in vivo*. Furthermore, the galactose-enriched cultures were highly sensitive to metabolically active pharmacological agents known to modulate the OXPHOS system, such as menadione, rotenone, resveratrol and metformin but no effect was observed with the glycolytic irreversible inhibitor, 2-deoxy-glucose. To build on these previously published results, we now report on a series of new studies designed to compare, for the first time, the transcriptional profiles using next-generation sequencing (NGS) of 3 differing pancreatic cell lines cultured in 2D versus 3D conditions, as well as with PDAC cells transplanted as xenografts *in vivo*. Our findings demonstrate the potential of genotypic analysis to better understand the perturbations of the OXPHOS system in a more disease-relevant 3D culture systems at the transcriptomic level. This approach may well be applicable to a wide range of therapeutic screening strategies, enabling the discovery of potential new and more effective treatments for PDAC.

## Materials and methods

### Materials

The LifeGel Digestion Kit and the LifeGel 3D cell culture plates for cancer spheroid formation (48-well) were both provided Real Research, Poland. Cell culture media were purchased from Capricorn Scientific GmbH, Germany (Dulbecco’s Modified Eagle Medium or DMEM with high glucose, L-glutamine and sodium pyruvate) or from Gibco, Europe (Iscove’s Modified Dulbecco’s Medium or IMDM). Cell culture plastics were from VWR International, Poland, unless otherwise stated. Fetal Bovine Serum or FBS was purchased from EURx, Poland and Tryp-LE from Santa Cruz Biotechnology, Germany. All 3 cell lines (Panc-1, CFPAC-1, MiaPaCa-2) were obtained from the European Collection of Authenticated Cell Cultures (ECACC), UK.

### 2D cell culture methods

2D cultures were first established for all 3 cell lines in conventional treated plastic flasks using either DMEM with 10% FBS (for Panc-1 and MiaPaCa-2) or IMDM with 10% FBS (for CFPAC-1). These methods facilitated the production of master stocks low passage numbers, which were then cryopreserved. Triplicate 2D cultures of each cell line were initiated from those master stocks and maintained in exponential growth until sufficient cells were available for harvesting and/or for 3D seeding. All experiments reported here were performed ensuring a difference of no more than 3 cell passages between the cultures used for analysis. 2D cultures were harvested by trypsinization, the cells counted and then prepared as a suspension in the appropriate pre-warmed culture medium. For each of the three human pancreatic cancer cell lines used in this study, 2D cultures prepared above were used in four separate ways: (1) for direct gene expression profiling as 2D cultures; (2) for injection into mice to produce cell-derived xenografts (CDX); (3) for seeding ‘young’ 3D cell cultures; or (4) for seeding ‘old’ 3D cell cultures. For the *in vivo* work, 0.5 x 10^6^ 2D cells were used for animal inoculations, having first been harvested, washed in phosphate-buffered saline (PBS) then resuspended in cell culture medium prior to ambient shipment to the Maria Sklodowska-Curie National Research Institute of Oncology (MSCI) in Poland. ‘Young’ 3D cell cultures were seeded with 10×10^3^ MiaPaCa-2 or Panc-1 cells or with 30×10^3^ CFPAC-1 cells per well; ‘old’ MiaPaCa-2 and Panc-1 3D cell cultures were seeded at the same density as for ‘young’, but ‘old’ CFPAC-1 3D cultures were seeded with 20×10^3^ cells per well. 32 wells were seeded for each of the three replicates for each experiment.

### 3D cell culture methods

LifeGel 3D cell culture plates (supplied in DMEM or IMDM medium) were pre-equilibrated at 37°C, which was gassed with 5% CO_2_. Medium was removed from above the LifeGel layer before seeding with cells (as described above) in 300µl of culture medium per well, after which plates were returned to the humidified incubator. The same culture media were used for 3D as for 2D with respect to each cell line. ‘Old’ 3D cell cultures were grown for a total of 14 days, with medium being replaced on days 6, 8, 10, 12 and 13. ‘Young’ 3D cell cultures were seeded with 10×10^3^ MiaPaCa-2 or Panc-1 cells or with 30×10^3^ CFPAC-1 cells per well; ‘old’ MiaPaCa-2 and Panc-1 3D cell cultures were seeded at the same density as for ‘young’. However, ‘old’ CFPAC-1 3D cultures were seeded with 20×10^3^ cells per well. 32 wells were seeded for each of the three replicates for each experiment. ‘Young’ 3D cell cultures were grown for 6 days without changing the medium before being harvested.

### 3D Cell harvesting

Cell-containing LifeGel was removed from plate wells along with medium using a 1ml pipette tip and pooled in a 50ml tube for each replicate. After centrifugation at 300 x g for 6 minutes, some of the medium-containing supernatant could be carefully removed to avoid disturbing the LifeGel-cell mixture. A 0.4 ml aliquot of LifeGel Digestion Kit was added to each 5ml of the mixture, before incubating for 30 minutes at 37°C with repeated mixing every 10 minutes. The digestion mix was then re-centrifuged, the supernatant carefully removed and cellular structures resuspended in PBS for cell number estimation. ‘Young’ 3D samples destined for animal inoculation were then repelleted and resuspended in a small volume of cell culture medium for transportation at ambient temperature to the MSCI.

### *In vivo* xenograft transplantation

All animals used for the CDX work were handled in strict accordance with good animal practice as defined by the relevant national and/or local animal welfare bodies. The animal work described in this study was approved by the Second Local Ethical Committee for Animal Research in Warsaw (permission no: WAW2/072/2022, dated June 8, 2022), which included adopting the relevant humane endpoint criteria: loss of weight, tumor volume (see below) and/or any adverse changes in mouse behavior that could be observed. Triple transgenic NSG-SGM3 immunodeficient mice that allow for the effective engraftment of human cells were used. The mice used were 13-28 weeks old and 29 female or male mice were implanted with the 3 different cell lines cultured under 2D or 3D conditions described above: MiaPaCa-2, Panc-1, CFPAC-1. To induce subcutaneous xenografts from the cultured cell lines, 0.3 - 1 × 10^6^ cells were injected subcutaneously (as a volume of 200µl) into the flank of each animal. The condition of all the mice were monitored daily for the relevant humane endpoint criteria described above. Tumor diameters were measured weekly with a caliper and their volume was calculated using the following formula: (length × width × width)/2. When tumors reached a volume of 1,500 mm^3^ or the mice reached an ethically determined humane endpoint not due to tumor size but based on observations of their overall condition as described above, the animals were humanely euthanized. Mice were allowed survive up to a maximum of 14 weeks. A necropsy was then performed, whereby the tumors were divided and collected. One mouse was found to be dead due to an unidentified reason on day 3 after implantation but had not appeared to be in any distress on the previous day. A sample of the tumor was snap frozen, stored at −80°C and preserved in 40% formalin before being transferred to 70% ethanol after 24 hours.

### Gene expression analysis

All the cellular samples described above were given an additional wash in PBS and aspirated pellets frozen at –80°C. CDX tissue samples comprising 2 mm cubes were frozen at –80°C after excision. Total RNA extraction, cDNA library preparations and whole transcriptome sequencing were performed at CeGat GmbH, using paired-end sequencing (2×100 bp) protocols and the Illumina NovaSeq platform.

### Bioinformatics analysis

Nucleotide sequences were stored in a FASTQ format before being trimmed and quality controlled using FASTQ software for further analysis [18]□. Gene counts were generated using Salmon software against concatenated transcriptomes (gencode: human v44 and mouse v.M33 including coding and non-coding transcripts) [19,20]□. Further analysis was performed in R environment utilizing DESeq2 tools and custom scripts, including heat-maps, principal component analysis, Venn diagrams, gene ontology analysis [21,22]□. Only transcripts with at least 5 reads in at least 3 samples were considered suitable for further analysis. A two-fold change and FDR <0.05 were required for differential gene expression change to be significant. Pathway similarity scores were calculated as fraction or percentage of genes changing in the same direction *in vivo* (from 3D) and 3D ‘old’ compared with 2D cultures. An average change in read counts was required to pass the 1.5-fold threshold for *in vivo* and 3D in comparison with the 2D cultures to be considered significant.

## Results

### The growth, morphology and reproducibility of cells in 2D and 3D cultures

To evaluate differences in morphology between the 2D and 3D cultures, triplicate 2D cultures of each cell line were seeded from master stocks and maintained in exponential growth until sufficient cells were available for harvesting and/or subsequent 3D seeding. All experiments were performed ensuring a difference of no more than 3 cell passages between them. Aliquots of LifeGel were added to each well of the tissue culture plates to provide a suitably thick layer of LifeGel. Aliquots of 100µl were added to each well in 96-well plates and 250µl in 48-well plates. The cells were then seeded on the surface of the LifeGel where they form 3D structures in the protein-based hydrogel, which is growth factor free. The cells then grew in multiple layers, migrating within the hydrogel and interacting with each other to create spheroid-like structures. The culture conditions recreate a cellular environment much closer to that typically encountered by a tumor growing in vivo. In this study, 6-day cultures were designated as being ’young’ 3D cultures and 14-day cultures were designated as ’old’ 3D cultures. A range of 3 cell lines were used in this study: CFPAC-1 from a ductal pancreatic adenocarcinoma derived by differential trypsinization of explant cultures, MIAPaCa-2 derived directly from pancreatic tumor tissue and PANC-1, which was isolated from a pancreatic carcinoma of ductal cell origin.

As can be seen from Fig 2, the images show distinct morphological differences between the 2D cultures, as well as between the 3D cultures. The morphologies of 2D monolayer cell cultures were as previously described for those cell lines and nothing unexpected was observed. ‘Young’ 3D cultures already clearly demonstrated cells growing at depth on top of each other and aggregating into distinctive 3D structures. CFPAC1 cells rapidly formed variously sized spheroidal structures with relatively smooth (and hence often ‘shiny looking’) outer surfaces. Once formed, these structures then became very slow growing. MiaPaca2 cells congregated or assembled into more loosely packed structures, resembling ‘large bunches of grapes’ in their appearance. Cells continue to divide rapidly in these more diffuse but expanding structures. PANC1 cell growth was less rapid than for MiaPaca2 cells and they formed predominantly discrete but ‘rough-surfaced’ spheroidal structures of different sizes that continued to enlarge within the hydrogel matrix.

**Fig 2.**
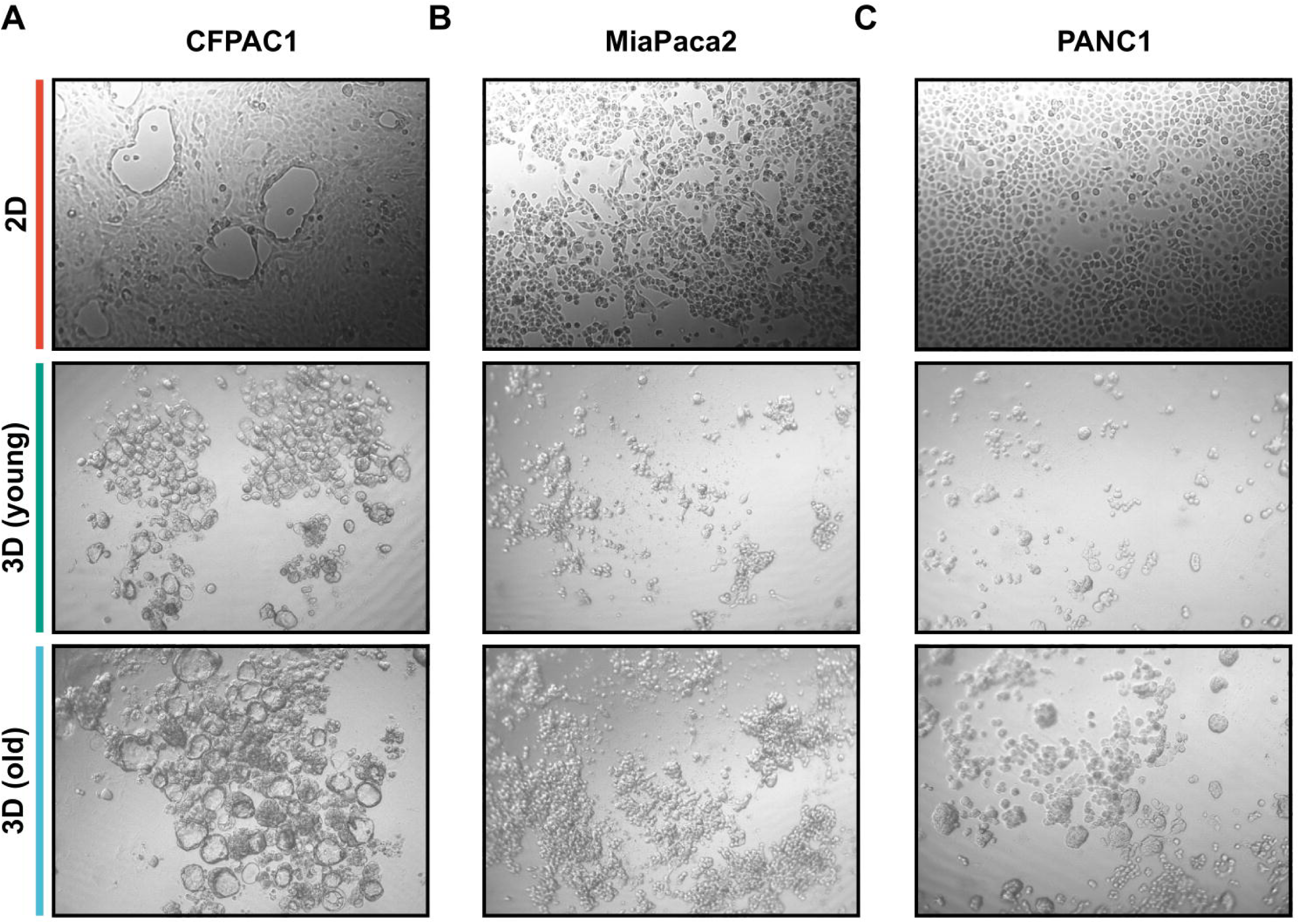
Generation and establishment of tumor-like spheroids. Microscopic image examples of 2D and 3D pancreatic cell line cultures (50x magnification, Zeiss Telaval 31, phase contrast 2D monolayer images, brightfield 3D images). (A) Derived from the CFPAC1 cell line. (B) Derived from the MiaPaca2 cell line. (C) Derived from the PANC1 cell line. 3D cell culture provides a reproducible platform for growth of cancer cell lines.

A key objective of the study was to demonstrate the reproducibility and consistency of the 3D systems *in vitro* and *in vivo*. To study mitochondrial oxidative phosphorylation genotype in these cultures at a transcriptomic level, therefore, we needed to establish controls for any significant variability across the differing culture platforms. Cultures were grown in standard 2D culture or 3D matrix, either for short or extended periods: 3D ‘young’ or 3D ‘old’. LifeGel was found to be reproducible batch to batch with a defined hydrogel stiffness and is compatible with standard imaging platforms, such as those used here. We decided to characterize the expression profiles using RNA-sequencing as a particularly sensitive method of analysis of the transcriptome. We observed robust reproducibility for representative genes: CD44 and Integrin Alpha Chain V (ITGAV) for the CFPAC1, MiaPaca2 and PANC1 cell lines (Fig 3A-C). CD44 is a cell-surface glycoprotein involved in cell–cell interactions, cell adhesion and migration. ITGAV is an integrin involved in adhesion and facilitating signal transduction. The expression levels of both genes are, therefore, sensitive biomarkers to test for reproducibility.

**Fig 3.**
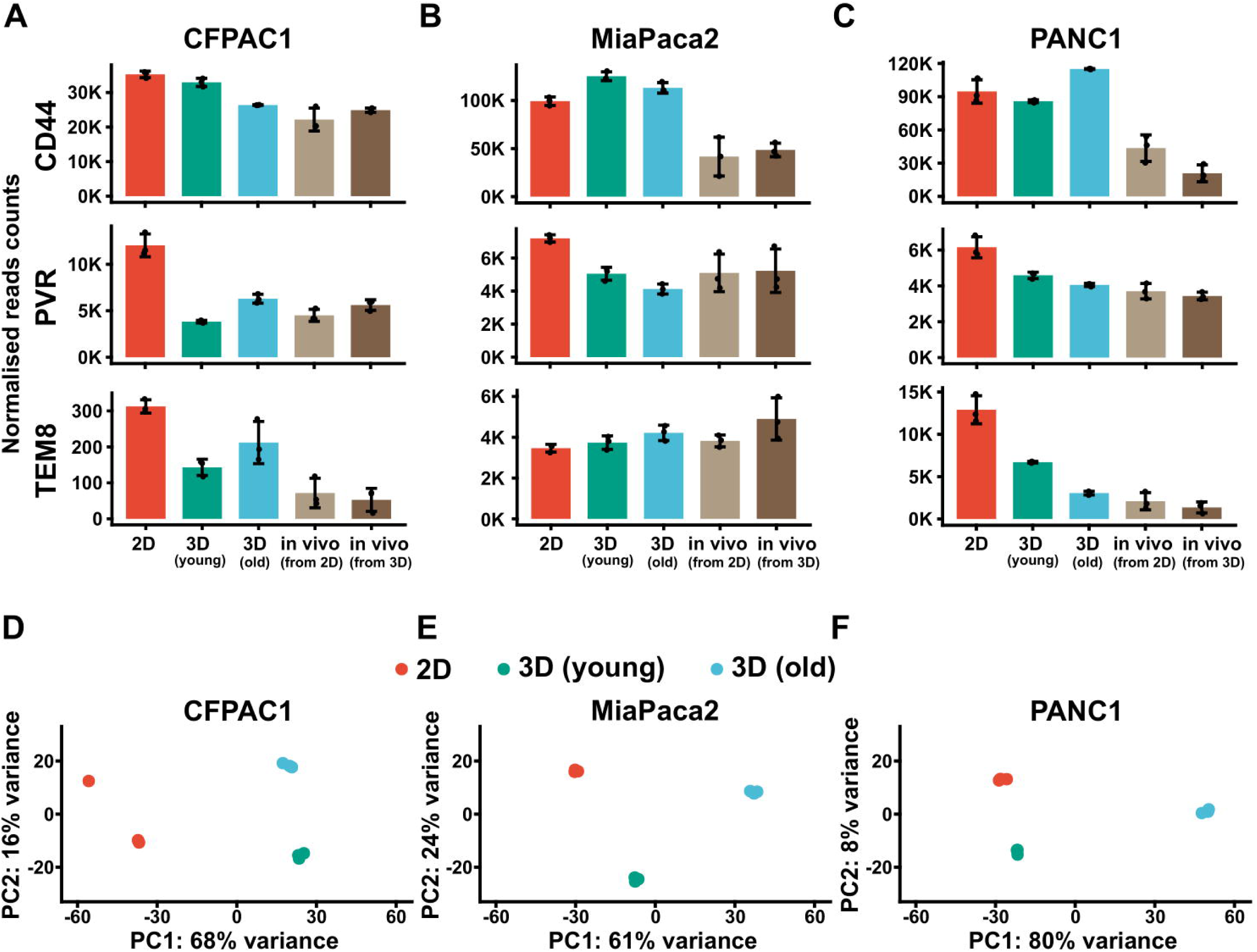
LifeGel 3D cell culture provides a reproducible platform for growth of cancer cell lines. (A-C) RNA-sequencing data results (normalized reads counts) for CD44, PVR and TEM8 genes for CFPAC1, MiaPaca2 and PANC1 cell lines (n=3). Cultures were grown in standard 2D culture or 3D matrix (either for short or extended period, 3D (young) or 3D (old). (D-E) Principal component analysis (PCA) for different transcriptome data summarizing expression profiles for distinct cell lines in 2D or 3D. Note the close clustering of the replicates, which is indicative of reproducibility.

Having established that expression profiles for selected genes were highly correlated between the different conditions, we then extended this analysis genome-wide. The global gene expression patterns by principal component analysis for all 3 cell lines whether grown in 2D or 3D showed close clustering of the replicates indicating high levels of experimental concordance across the culture platform (Fig 3D-F) to underpin the further studies described below.

### Commonalities in transcriptional trajectories between *in vivo* and 3D cultures

The relevance of the 3D *in vitro* culture conditions to the same cell lines grown *in vivo* is illustrated in Figure 4. The analysis compared the upregulated or down regulated genes with the baseline 2D data, which revealed that there is statistically significant overlap between the transcriptome of ‘old’ 3D cultures and the same cells grown in the *in vivo* CDX model. In our analysis, we considered both the coding and non-coding genes. To exemplify which processes are most affected *in vivo* and in 3D, we selected genes that were downregulated in these conditions compared with 2D cultures. Indeed, we observed significant correlations across metabolic processes that include respiration and/or mitochondrial functions. In addition, we observed that pathways that related to genome organization seem to be repressed in 3D cultures and *in vivo*, when compared with 2D cultures.

**Fig 4.**
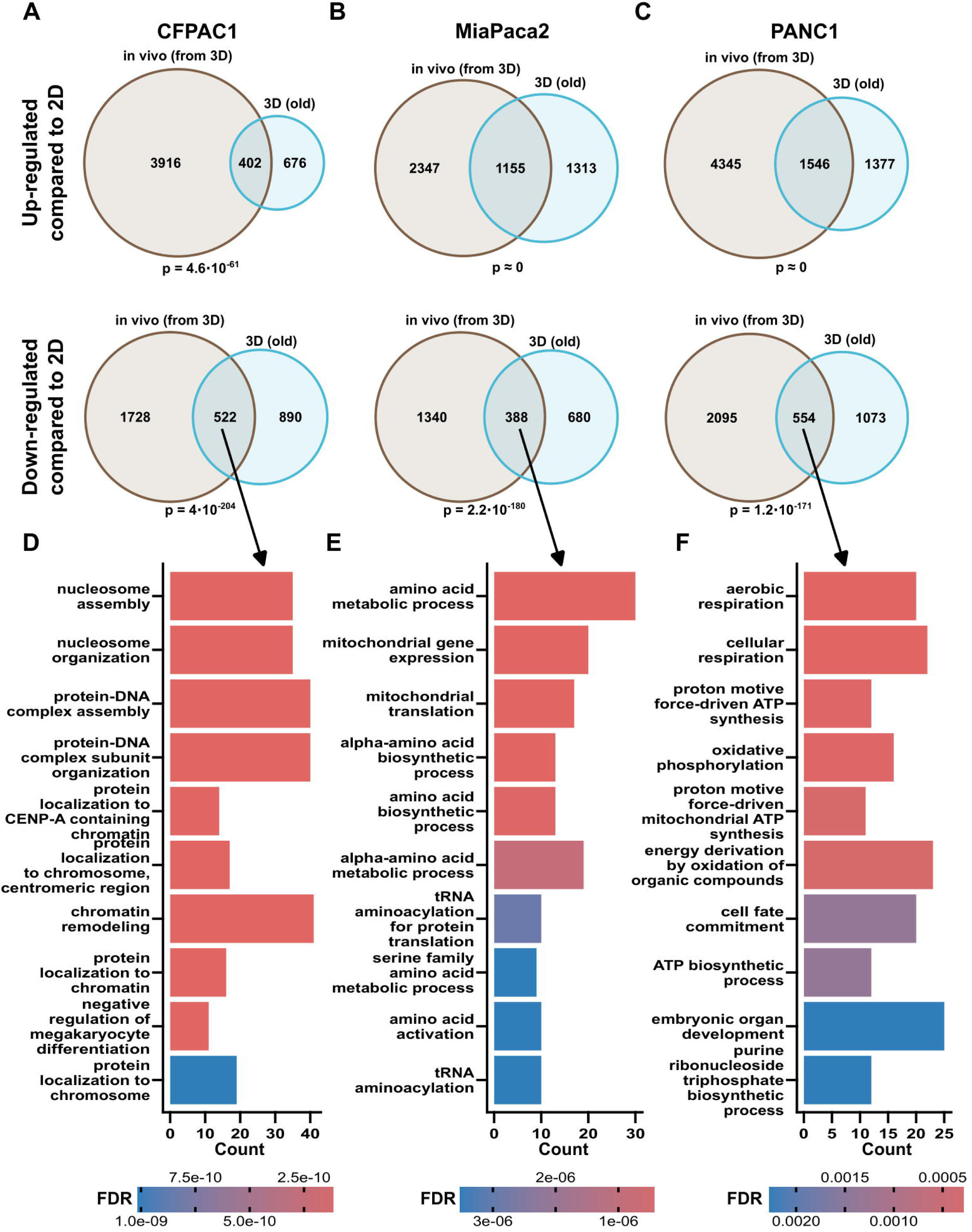
Commonalities in transcriptional trajectories between *in vivo* and 3D cultures. (A-C) Common genes significantly up-regulated or down-regulated *in vivo* (from 3D) or 3D (old) in reference to 2D cultures. Overlap between in vivo and 3D was tested using hypergeometric test indicating significant association. Both coding and non-coding genes were included. (D-F) Ten most enriched pathways for genes that are down-regulated both *in vivo* (derived from 3D) and 3D (old).

### Similarities between *in vivo* and 3D cultures at the OXPHOS pathway level

To further explore parallels in gene expression in 3D and the *in vivo* CDX models, we devised a pathway similarity score to count genes in any given pathway, if it followed the same trends in conditions in question. If gene was down- or upregulated both *in vivo* and in 3D, it was assumed to have changed. Although not every parameter correlated, OXPHOS-related genes score particularly strongly when using this method (Fig 5A-C). Additionally, we plotted genes from OXPHOS pathway as a heatmap to show that a substantial number of genes changed in both the *in vivo* and ‘old’ 3D cultures, when compared with 2D cultures (Fig 5D-F). OXPHOS-related genes were more affected *in vivo* and in 3D than they were in 2D cultures. The global trend observed was that 3D is better at mimicking the OXPHOS-related gene expression profile than were the 2D cultures.

**Fig 5.**
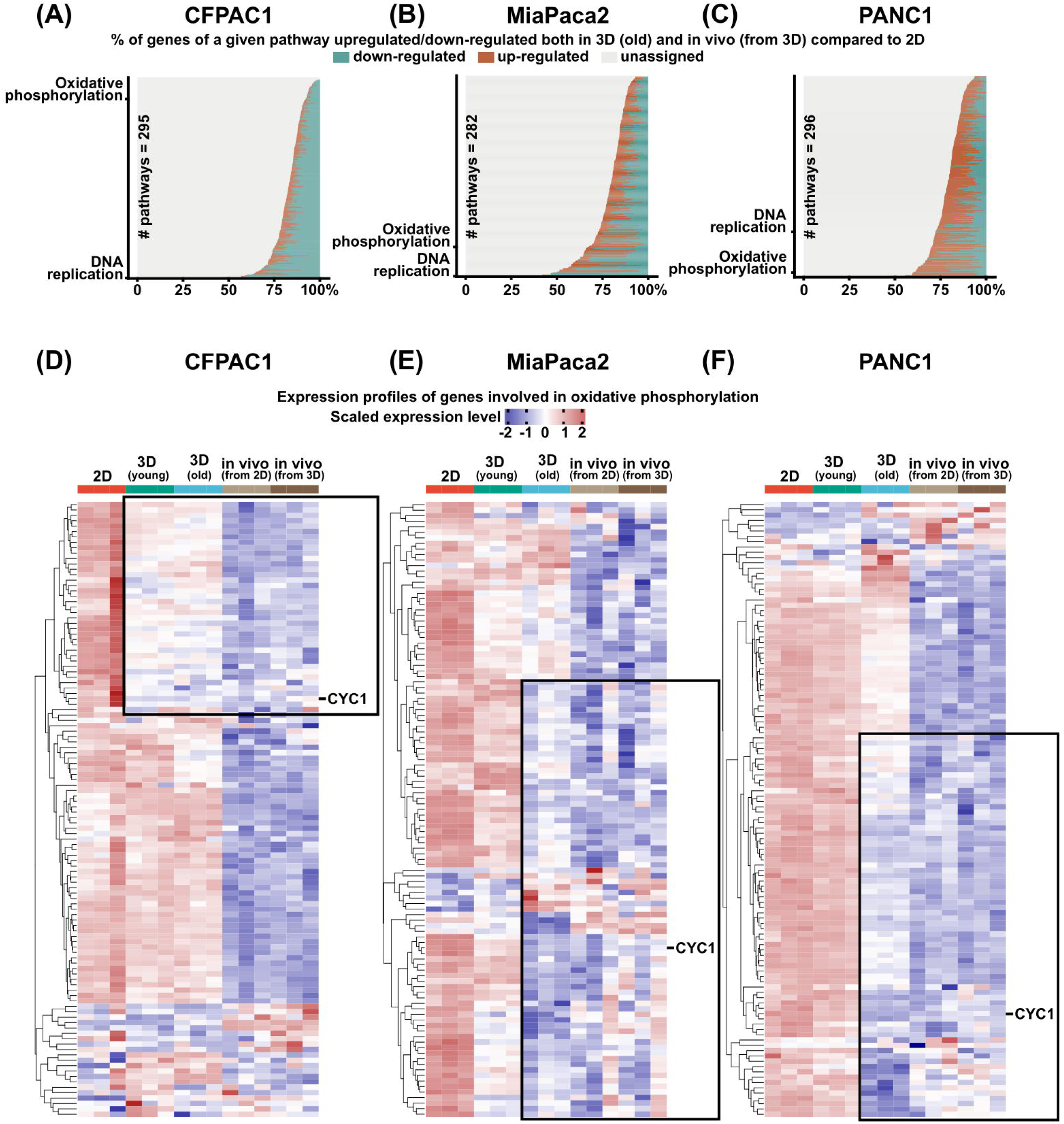
Similarities between *in vivo* and 3D at the pathway level. (A-C) Pathway similarities between the samples were evaluated as a percentage of genes in a given pathway that exhibited the same direction of change *in vivo* CDX and ‘old’ 3D culture samples when compared with 2D culture samples. Any gene in a pathway was considered to be up- or down-regulated only if the average gene count was at least 1.5-fold higher or lower both in 3D and *in vivo* when compared with 2D culture. When this criterion was not met, that gene was set as unassigned. The pathways were sorted according to the lowest percentage of unassigned genes for a given pathway. Pathways with at least 25 genes were considered and only two pathways are highlighted using this methodology: OXPHOS and DNA replication. (D-F) Heatmap depicting expression profiles of genes involved in oxidative phosphorylation (KEGG: hsa00190) among different cell lines and conditions. Only the Cyc1 gene is highlighted as a reference. Black rectangles encircle the cluster of genes that show similarities between 3D cultures and *in vivo* samples.

## Discussion

3D cultures represent an attractive screening assay platform to capture disease-associated pathways *in vitro* that are not so well represented in conventional 2D cultures. Here, i*n vitro* 3D culture of pancreatic cancer cells resembled the aberrant mitochondrial oxidative phosphorylation genotype observed *in vivo* when analyzed at a transcriptomic level.

It is important to understand how these findings can fit within the context of drug discovery. Discovering new drugs is a costly, lengthy and largely unpredictable endeavor. Overall, the number of new drugs approved per billion dollars spent on research and development has halved about every nine years from 1950 to 2010 with no obvious signs of any substantial improvements over the subsequent decade [11].

Previous analyzes have uncovered how assay validity and reproducibility can be correlated across a range of screening assays and disease models, such as in the key publication by Scannell and Bosley in 2016 [23]. These authors proposed that increasing the implementation of more relevant and predictive screens should be incorporated pre-clinically into the drug discovery process. A more rigorous understanding of efficacy and toxicity at multiple biological levels would then offer a potential solution to this systemic productivity problem. It has been proposed that pivotal decision-making assays, such as those described here, need to be introduced much earlier into the discovery process to enable disruptive changes in drug discovery to make a real difference to productivity [13]. Analytical methods based on decision theory have demonstrated that slight changes in the ’predictive validity’ of an assay can have a remarkably significant impact on downstream success rates [11].

3D cell cultures, such as those described in this study, are typically derived from cell lines that have previously been grown in monolayers or suspensions and then aggregated into 3D-compatible culture systems. If grown in suitable hydrogels, spheroids allow cells to communicate with each other in an analogous manner to an *in vivo* 3D environment and can mimic tumors if derived from appropriate cell lines. Therefore, evaluating and then improving the predictive potential of such pre-clinical assays *in vitro* could be used to better predict future outcomes in human clinical trials. Novel screening strategies could then translate the medical benefits to cancer patients.

More specifically, in relation to pancreatic cancer, metabolic phenotypes can be transient [24]. Progressively de-differentiated pancreatic cancer cells have been shown to undergo a shift from glycolytic to oxidative metabolism, resulting in a quiescent state. Following the re-differentiation, these cells regained their proliferative capacity and glycolytic metabolism, commonly associated with a greater aggressiveness. Metabolic switches in PDAC cells can occur as part of the underlying pathology but also as a response to disruptions caused by treatments. Therefore, it is both timely and opportune to better understand the underlying OXPHOS mechanisms in PDAC, as modulating these pathways therapeutically could help either to prevent or overcome the metabolic plasticity in patients. Clearly, considerable metabolic plasticity is observed across many wide-ranging biochemical pathways in cancers, including the well-characterized Warburg effect, but PDAC tumors could be more specifically targeted to mitigate or even reverse the aberrant effects of the metabolic OXPHOS phenotype described here. This hypothesis is illustrated in Fig 6.

**Fig 6:**
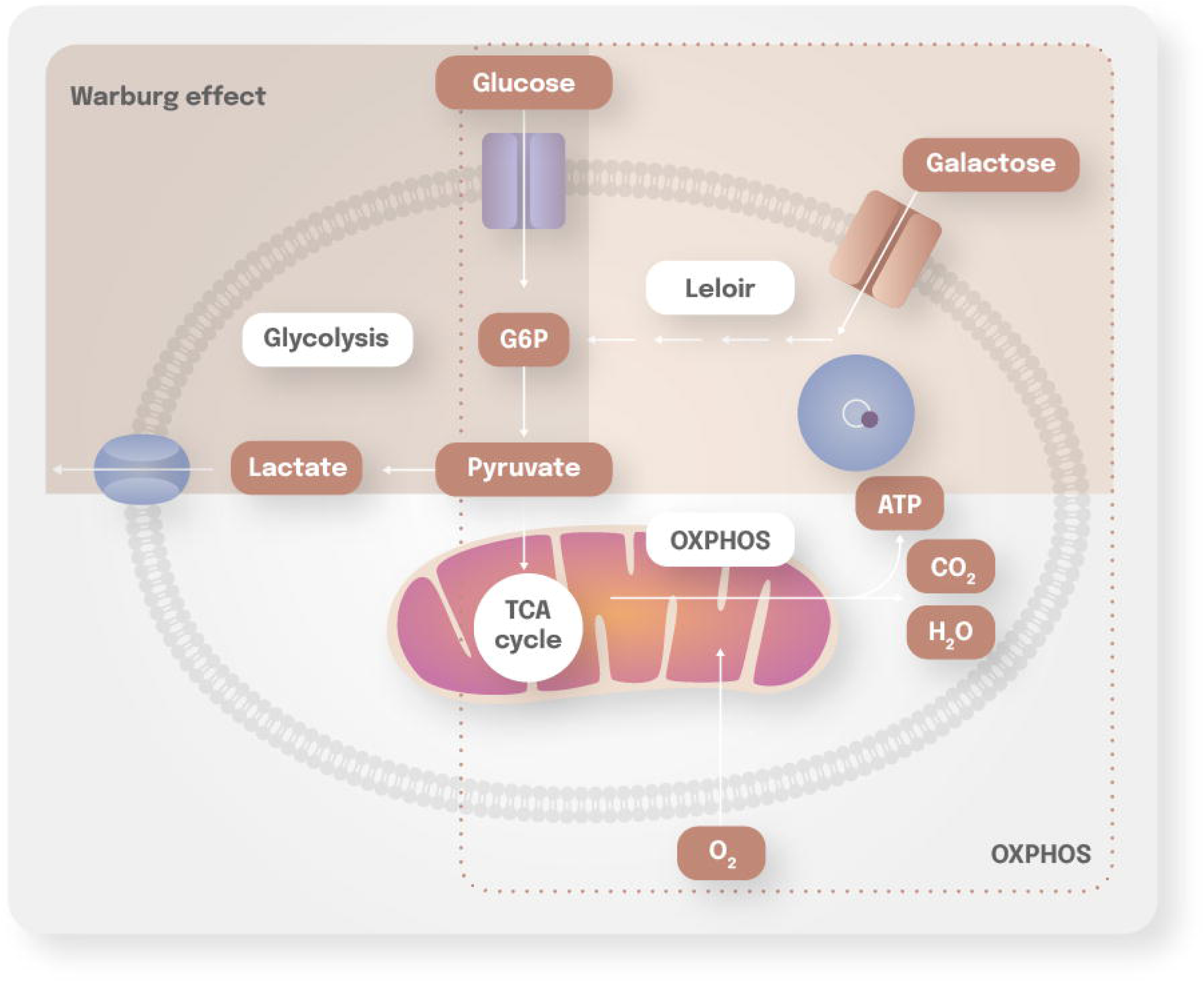
Schematic representation of Warburg and OXPHOS metabolism. The Warburg effect is based on the observation that many cancer cells release energy predominantly through the inefficient process of aerobic glycolysis (and named after Otto Warburg). The Leloir pathway is a metabolic pathway for the catabolism of D-galactose (named after Luis Leloir). The alternative OXPHOS in the mitochondria is also illustrated for comparison.

For the OXPHOS pathway to be further evaluated as an emerging target pathway for new cancer therapies [25], then 3D cultures relevant to PDAC need to be implemented. The 3D culture systems described in this study could very well become an important testbed for evaluating and triaging suitable pharmacological options in drug discovery, as well as for other therapeutic strategies.

## Supporting information

Supplemental image

## Acknowledgments

We acknowledge Sharla White, Hannah Buck and Amy Lam for their help in producing the video in the supporting information.

## Abbreviations

2D: 2 dimensional
3D: 3 dimensional
ATP: Adenosine triphosphate
CDX: cell-derived xenografts
DMEM: Dulbecco’s Modified Eagle Medium
ECACC: European Collection of Authenticated Cell Cultures
FBS: Fetal Bovine Serum
IMDM: Iscove’s Modified Dulbecco’s Medium
MSCI: Maria Sklodowska-Curie National Research Institute
NGS: next-generation sequencing
OXPHOS: oxidative phosphorylation
PBS: phosphate-buffered saline
PDAC: pancreatic ductal adenocarcinoma
RNA: ribonucleic acid.

## Supporting information

S1. Video showing 3D imaging of PANC-1 cell cultures.

