## Supplementary figures and images for "3D culture of pancreatic cancer cells *in vitro* recapitulates an aberrant mitochondrial oxidative phosphorylation genotype observed *in vivo*"

### Supplemental image

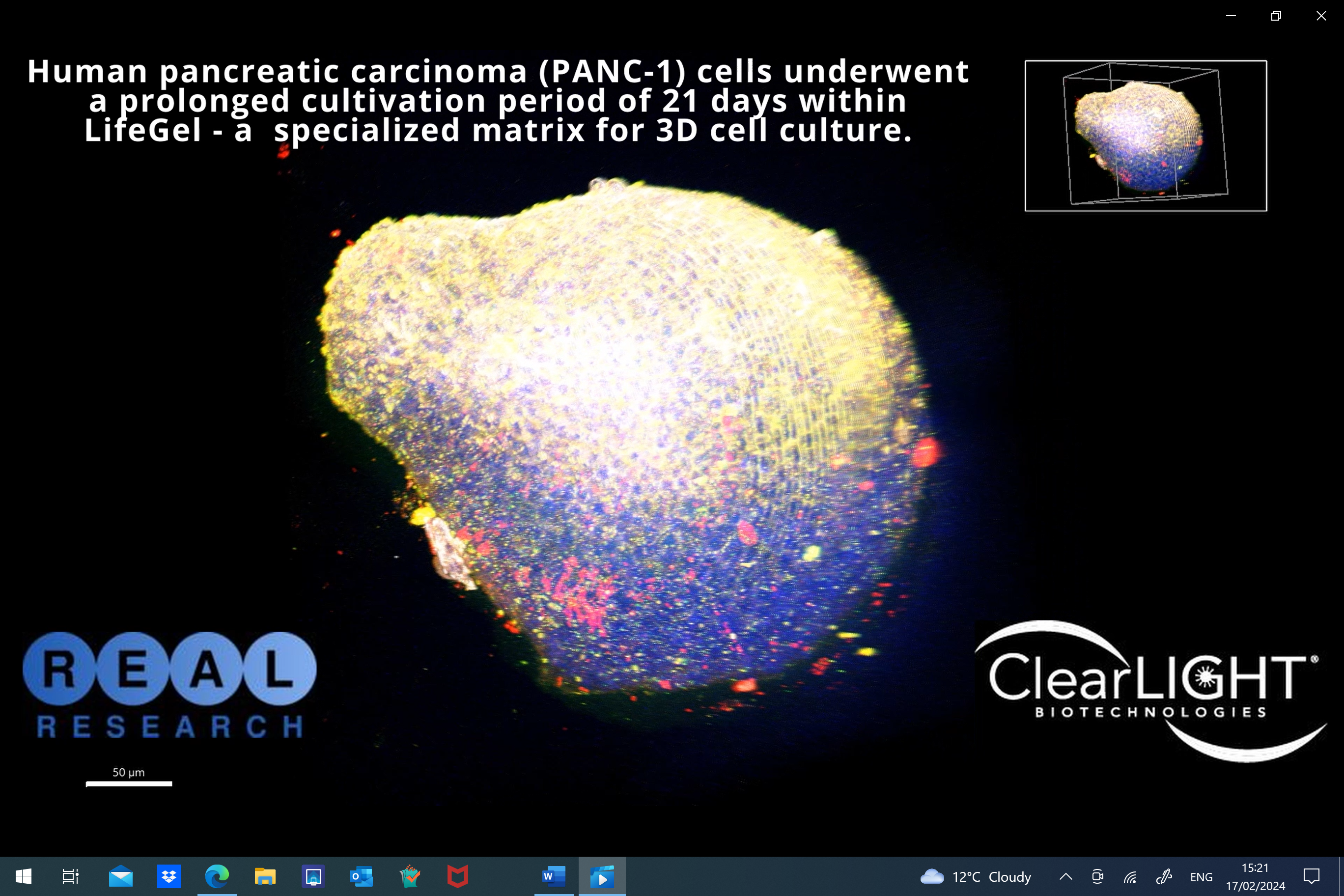
